# An intrinsic endothelial dysfunction causes cerebral small vessel disease

**DOI:** 10.1101/2021.12.13.472377

**Authors:** Sophie Quick, Tessa V. Procter, Jonathan Moss, Angus Lawson, Serena Baker, Marc Walton, Mehreen Mohammad, Will Mungall, Ami Onishi, Zuzanna Tobola, Michael Stringer, Maurits A. Jansen, Antoine Vallatos, Ylenia Giarratano, Miguel O. Bernabeu, Joanna M. Wardlaw, Anna C. Williams

## Abstract

Small Vessel Disease (SVD) is the leading cause of vascular dementia, causes a quarter of strokes, and worsens stroke outcomes(1, 2). The disease is characterised by cerebral small vessel and white matter pathology, but the underlying mechanisms are poorly understood. Classically, the microvascular and tissue damage has been considered secondary to extrinsic factors, such as hypertension, consisting of microvessel stiffening, impaired vasoreactivity and blood-brain barrier dysfunction identified in human sporadic SVDs. However, increasing evidence points to an underlying vulnerability to SVD-related brain damage, not just extrinsic factors. Here, in a novel normotensive transgenic rat model where the phospholipase flippase *Atp11b* is deleted, we show pathological, imaging and behavioural changes typical of those in human sporadic SVD, but that occur without hypertension. These changes are due to an intrinsic endothelial cell dysfunction, identified in vessels of the brain white matter and the retina, with pathological evidence of vasoreactivity and blood-brain barrier deficits, which precipitate a secondary maturation block in oligodendroglia and myelin disruption around the small vessels. This highlights that an intrinsic endothelial dysfunction may underlie vulnerability to human sporadic SVD, providing alternative therapeutic targets to prevent a major cause of stroke and dementia.

## Introduction

Small vessel disease (SVD) causes about 45% of dementias, including most vascular dementia and cognitive impairment, as well as 25% of ischemic stroke and most haemorrhagic strokes in older people(2, 3), making it a major target in tackling the global burden of neurodegenerative disease. SVD affects the small perforating blood vessels of the brain and is diagnosed using a collection of features on magnetic resonance imaging (MRI), including white matter hyperintensities (WMH), lacunes and microbleeds(3), with the increased extent of these white matter changes correlating with worse cognition(4, 5). In addition to presenting with stroke or cognitive impairment, SVD may also cause impaired gait or balance, and neuropsychiatric symptoms including depression. However, the clinical expression is often ‘covert’ and under-recognised both by affected individuals and clinicians(6–9). Indeed, studies of the general population show that features of SVD are common, increase in prevalence with age and are associated with hypertension, diabetes and hypercholesterolaemia(10). SVD is often considered to result from hypertension(11), other vascular risk factors, or atherothromboembolic disease as in most large artery strokes. While hypertension is certainly a risk factor for developing SVD(12), approximately 30% of patients with sporadic SVD have no history of hypertension(13). Additionally, all common vascular risk factors combined only account for a small percentage of variance in WMH severity(14) and, consistent with this, it has been difficult to demonstrate that anti-hypertensive treatment can prevent worsening of MRI or clinical features(12).

This suggests that the risk of developing SVD is not simply a consequence of exposure to vascular risk factors in adulthood. Indeed, our epidemiological data show that increased SVD severity in later life associates with early life factors such as lower cognitive ability in youth (and subsequent lower educational attainment)(15), independent of adult risk factor exposure(16). Since cognitive ability is partly explained by white matter integrity(17), the association with SVD in later life may reflect inherently poorer white matter integrity from youth, and hence white matter vulnerability to damage(3). Consistent with this, the integrity of normal appearing white matter is reduced in young adults who have an increased genetic risk for WMH, long before WMH develop(18).

The link between early life factors/white matter changes and vascular disease may lie with the endothelial cell of the blood-brain barrier. Using the inbred Spontaneously Hypertensive Rat - Stroke Prone strain (SHRSP), a well-established rat model with many features of sporadic human SVD(19–21), we previously demonstrated intrinsically dysfunctional ECs with reduced endothelial nitric oxide synthase (eNOS) and secondary white matter changes in young rats prior to onset of hypertension, due to block of oligodendroglial maturation through increased EC secretion of Heat shock protein 90 alpha (HSP90α)(22). As these changes were also present in *ex vivo* SHRSP brain slices from postnatal rat pups after 3 weeks of culture (where there is no blood flow or pressure), this confirmed EC dysfunction was intrinsic, and that hypertension is not the sole cause of the SVD. By comparing the SHRSP whole genome sequence to that of its closest relative the Spontaneous Hypertensive Rat (SHR) (which is hypertensive without SVD changes), we identified a homozygous exonic deletion mutation in *Atp11b* in the SHRSP but not in the SHR. Loss or knock down of *Atp11b/ATP11B* in cultured rodent or human ECs was sufficient to mirror the SHRSP EC dysfunction and secondary OPC maturation block. We also identified a SNP in *ATP11B* that associated with human WMH in the CHARGE consortium(22). This combined evidence led to our hypothesis that hypertension is not required for SVD pathology in rat or human, which we test here in a novel transgenic *Atp11b*KO rat model.

## Results

### *Atp11b*KO rat generation

The SHRSP has a deletion mutation in *Atp11b* which predicts a truncated ATP11B protein, but leads to its total loss by western blot with an N-terminal antibody(22). With the company Cyagen, we reproduced this deletion mutation in a new transgenic rat model on a Sprague-Dawley background, using CRISPR-Cas9 technology, with guide RNAs which disrupt from exon 5 to exon 8 (**Fig. 1A and methods**). Loss of ATP11B protein was confirmed with an ATP11B antibody by immunofluorescence (**Fig. 1B**). These rats breed normally and reached 1 year of age without overt health problems.

**Fig. 1.**
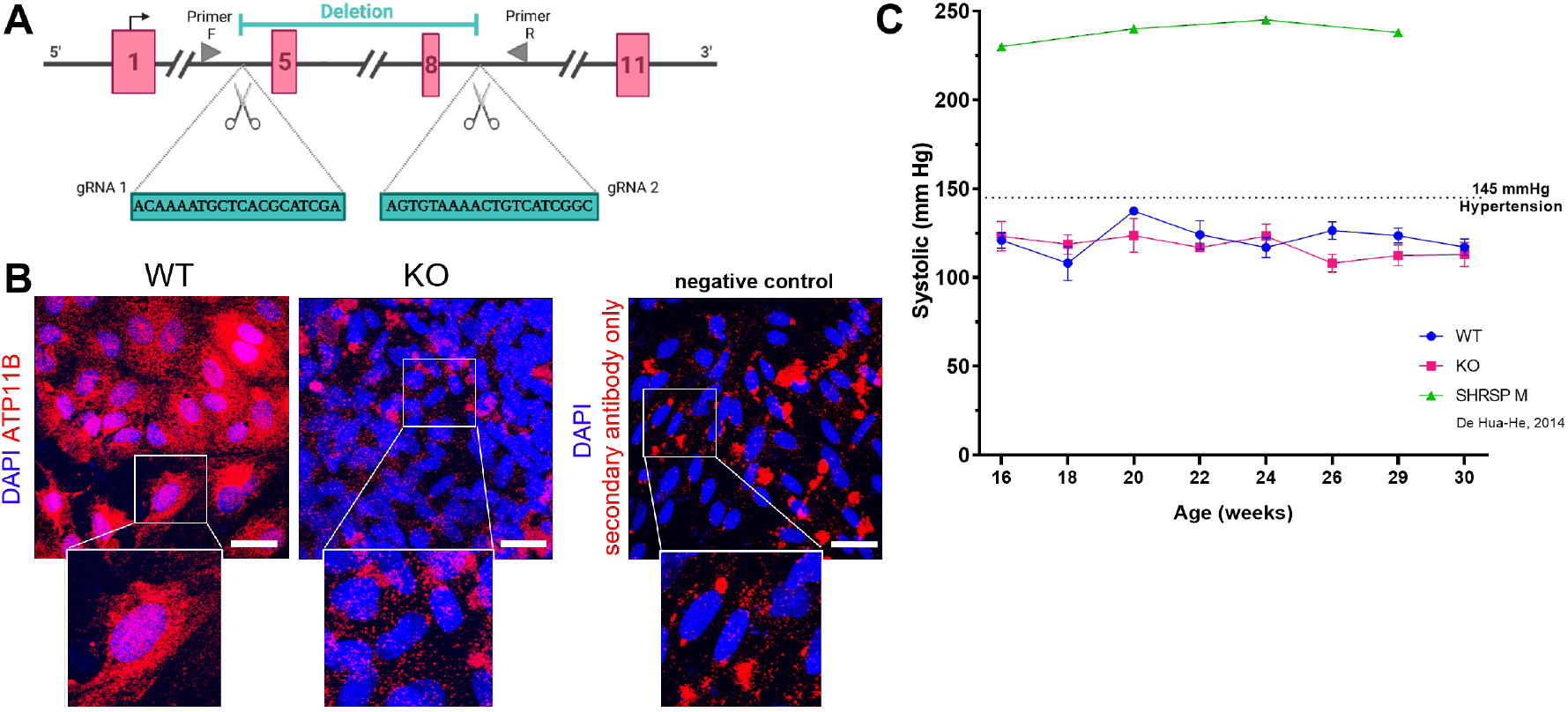
*Atp11b*KO rat is normotensive. **(A)** *Atp11b*KO CRISPRCas9 deletion strategy. gRNAs induce NHEJ leading to deletion between exons 5 and 8, with knockout confirmed by PCR using primers in locations indicated. **(B)** Immunostaining for ATP11B on cultured brain endothelial cells from WT and KO perinatal animals indicates a knockout at the protein level, with KO cells and secondary antibody only showing background staining. **(C)** Longitudinal blood pressure study on two groups of mixed-sex KO (11 animals) and age-matched controls (15 animals) shows that KO animals are normotensive (>145mmHg denoted as hypertension), and do not diverge from WT, unlike hypertensive male SHRSPs (data from De Hua-He et al., 2014). Graph shows mean +/-SD for whole group, using the average systolic reading from each animal.

### The *Atp11bKO* rat is normotensive

Our strategy of selecting a mutation found in the SHRSP, a rat which has SVD pathology, but not found in the SHR, a rat with hypertension but lacking overt SVD pathology, led to our prediction that the *Atp11b*KO rat would be normotensive. In a longitudinal study, there was no difference in blood pressure of the *Atp11b*KO rat compared to wildtype (WT) (Sprague-Dawley) controls at any age, with the average reading not surpassing 145mmHg in either group, denoted as hypertension in Sprague-Dawley rats(23) (**Fig. 1C**), and no difference between males and females. Next, we investigated whether, without hypertension, there were still signs of EC dysfunction at a juvenile age (3-6 weeks old) and an adult age (24-30 weeks old).

### *Atp11b*KO-ECs show signatures of dysfunction despite the absence of hypertension

EC dysfunction can be defined by several molecular signatures, including loss of eNOS indicating less NO production, reduction in Claudin 5 (CLDN5) tight junction protein indicating loss of BBB integrity and an upregulation in activation markers such as Intercellular Adhesion Molecule 1 (ICAM-1)(24–26), that parallel the microvascular dysfunctions observed macroscopically in human SVD. ECs cultured from the brains of neonatal *Atp11b*KO rats (KO-ECs) compared to WT-ECs show a significant reduction in the amount of eNOS (**Fig. 2A**), a significant reduction in amount of CLDN5 and a change in its location, as defined by western blot and immunofluorescence respectively (**Fig. 2B, C**) and a significant upregulation of ICAM-1 (**Fig. 2D**). This suggests early inherent EC dysfunction similar to that seen in 3 week old SHRSPs(22). As SVD increases with age, with MR changes typically in the periventricular and deep white matter, we also analysed markers of EC dysfunction in the deep frontal white matter of rat brain at the juvenile and adult ages. We found significantly fewer CLDN5+ blood vessels (Isolectin(Iso)B4+) overall in *Atp11b*KO juvenile rats (**Fig. 2E**). This reduction is less clear in the adults, as the amount of CLDN5+ blood vessels is more variable. Therefore, we focused on EC ultrastructural abnormalities in the same area, using electron microscopy (EM) analysis. Four blinded independent researchers graded EC pathology as 0 (normal), 1 (mild) and 2 (severe abnormality), finding increased EC inner surface undulations, luminal blood cell stasis and abluminal thickened basement membrane in adult KO animals compared to controls (**Fig. 2F, suppl. Fig.1**). These changes are more frequent in KO animals (**Fig. 2G**), indicating EC dysfunction at an ultrastructural level, consistent with structural abnormalities seen in small perforating brain blood vessels in human SVD(27). Not all blood vessels are affected, with heterogeneity between vessels even within animals (**Fig. 2H**), highlighting the well-recognised focal nature of SVD as found in humans(27).

**Fig. 2.**
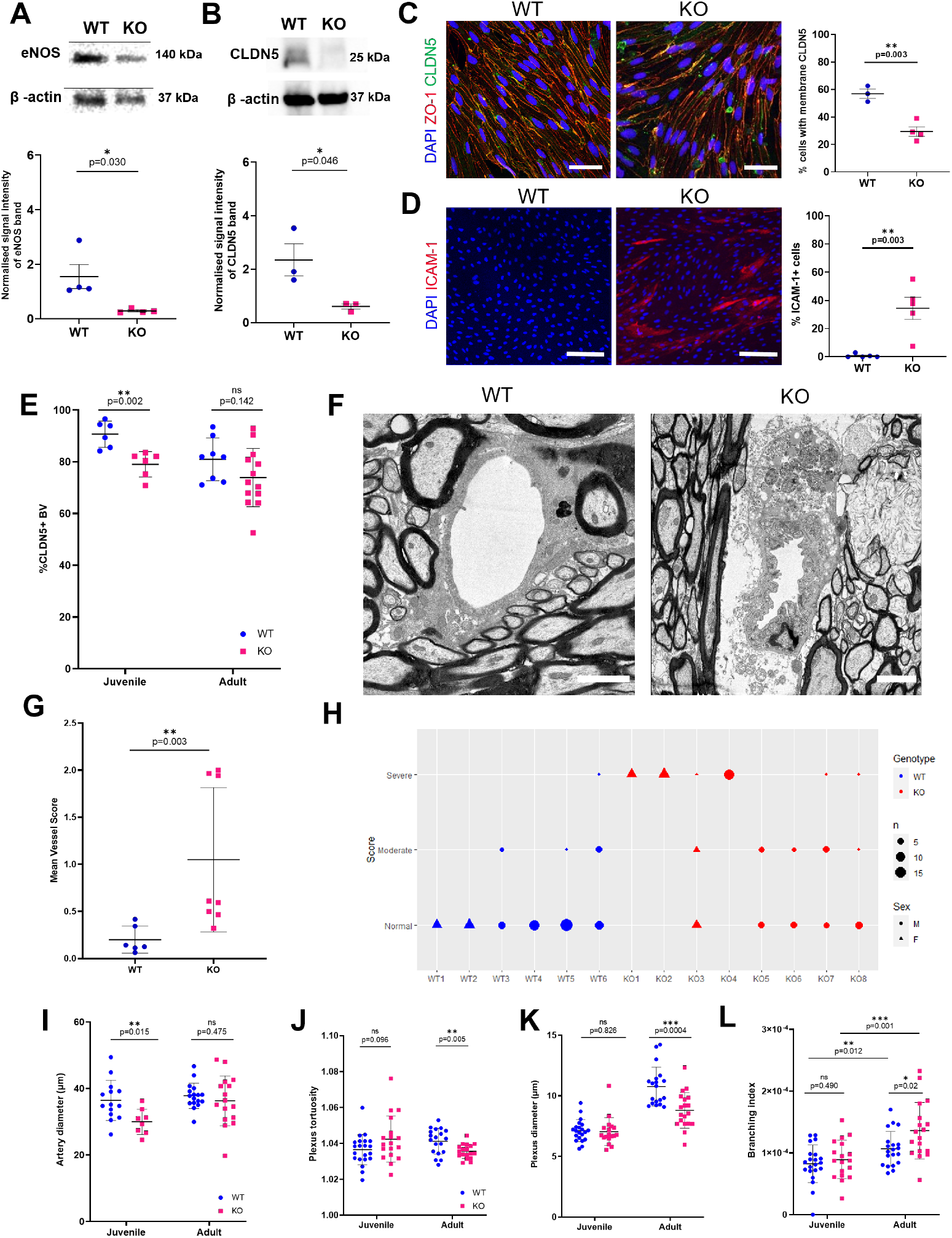
*Atp11b*KO-ECs are inherently dysfunctional. KO-ECs show dysfunction with expression of significantly less **(A)** eNOS (t-test, p=0.030, t=2.825, df=6) and **(B)** CLDN5 (t-test, p=0.046, t=2.826, df=4) than WT-ECs - western blot of cell lysates, with quantification by densitometry (relative to β-actin) below, calculated from images taken with LiCor image system for eNOS and from photographic film for CLDN5. **(C)** Fewer cultured KO-ECs express the tight junction marker CLDN5 (green) compared to WT-Ecs at the border between all neighbouring cells, delineated by ZO-1 (red), (DAPI-stained nuclei - blue) with quantification of % cells with membranous CLDN5, (t-test, p=0.003, t=5.553, df=5) scale bar = 20µm. **(D)** More cultured KO-ECs express the endothelial activation marker ICAM-1 compared to WT-ECs, with quantification (t-test, p=0.003, t=4.306 df=8), p=0.003, t=4.306 df=8). **(E)** Fewer tight junction CLDN5+ blood vessels in *Atp11b*KO tissue from juvenile age group compared to WT (t-test, p=0.002, t=4.018, df=10). **(F)** Marked abnormalities in adult KO blood vessels shown with electron microscopy (scale bars 2µm), **(G)** Abnormalities in ECs quantified as mean vessel score per animal (Mann-Whitney U =2, p=0.003), full descriptors used to score are found in supplementary methods. **(H)** Heterogeneity is seen between vessels within animals; plotted as modal rater scores for each vessel analysed. **(I)** Retinal artery diameter is significantly smaller in juvenile KO animals compared with WT animals (t-test, p=0.015, t=2.688, df=20) but not at the adult age (t-test p=0.475, t=0.724, df=31). Adult KO retinal plexus vessels show **(J)** significantly less tortuosity (t-test, p= 0.005, t=2.969, df=35), **(K)** significantly smaller diameter (t-test, p=0.0004, t=3.934, df=36), and **(L)** significantly increased branching (t-test, p=0.020, t=2.430, df=36), with increased branching with age in both groups: KO animals (t-test, p=0.001, t=3.584, df=34), WT animals (t-test, p=0.012, t=2.641, df=39). There are no identified differences between plexus measurements by genotypes at the juvenile age (t-tests, Diameter: p=0.826, t=0.221, df=37; Branching: p=0.490, t= 0.6974, df=37; Tortuosity p=0.096, t=1.710, df=37). All graphs show mean +/- SD.

We next examined the larger retinal arterioles/venules and smaller plexus vessels in flat mount retinal preparations(28) labelled with IsoB4 in juvenile and adult animals. The retina can be considered as a ‘window’ to the brain, with shared embryological origin and neurodegenerative pathways, and as retinal vessels are relatively easily measurable *in vivo*, retinal imaging is being explored as a non-invasive diagnostic tool for SVD in patients(29). A reduced retinal artery (but not venule) diameter was observed within the KO animals in juveniles only (**Fig. 2I, Suppl. Fig. 2**). Meanwhile, reduced diameter and tortuosity but increased branching of plexus vessels were noted exclusively in the adult animals (**Fig. 2J-L**). We speculate that this reflects early impaired vascular vasodilation (possibly secondary to NO loss) and a later compensatory response to hypoxia leading to increased plexus vessel branching via an angiogenic response, supported by the increase in branching index in adult KO animals compared to juveniles (**Fig. 2L**). Human fundus imaging studies show that reductions in retinal arteriolar diameter, density and fractal dimension are associated with SVD features on MRI, independent of vascular risk factors(30), and plexus vessels can be imaged using optical coherence tomography angiography. Although some of these human retinal features are present with hypertension, in these KO rats, we can be clear that these abnormalities are independent of hypertension.

Thus, hypertension is not required to produce EC dysfunction and blood vessel changes in the *Atp11b*KO rat. Next, we investigated whether white matter changes were also present.

### The *Atp11b*KO rat has white matter changes similar to those in human sporadic SVD

In the *Atp11b*KO brain deep frontal white matter, there were no gross general myelin defects, as assessed by Proteolipid protein (PLP) immunofluorescence, although there was the expected increase in amount of myelination between juveniles and adults as most myelination occurs postnatally (**Fig. 3A**). However, there were significantly fewer mature Olig2+NogoA+ oligodendroglia in the deep frontal white matter in the juveniles but not in the adults suggesting a WM disturbance (**Fig. 3B**). This shows a similar pattern to the reduction in CLDN5 expression in small blood vessels in the same area of juvenile brains only (**Fig. 2E**), suggesting that EC-secreted factors change in how they affect oligodendroglia over time, possibly reflecting some compensation. In human SVD, subtle diffuse WM changes are seen by MRI(31, 32), but pathological white matter changes are usually described as focal around blood vessels(27) and so we quantified the PLP-positive immunofluorescence pixels in a 10µm concentric ring around small vessels of the deep white matter, with the hypothesis that there would be less myelin nearer to blood vessels. Surprisingly, there was a significant increase in the number of PLP-positive pixels found near these vessels in the *Atp11b*KO adults (**Fig. 3C**). To explain this finding, we returned to EM to investigate the ultrastructure of the myelin in adult animals, hypothesising that the increased PLP labelling may be secondary either to PLP-positive myelin debris around vessels or less compact myelin sheaths. Using a pathological grading system of 0 (normal), 1 (mild) and 2 (severe abnormality), we found evidence of perivascular abnormalities around blood vessels (**Fig. 3D**), with increased myelin debris (yellow arrowhead), vacuoles and demyelinated axons, but no clear myelin compaction changes. These perivascular changes appeared mostly in KO animals, but with variability between vessels from the same animals and between animals (**Fig. 3E,F**) consistent with our PLP-immunofluorescence data around these vessels (**Fig. 3C**) which also showed this vessel-to-vessel heterogeneity.

**Fig. 3.**
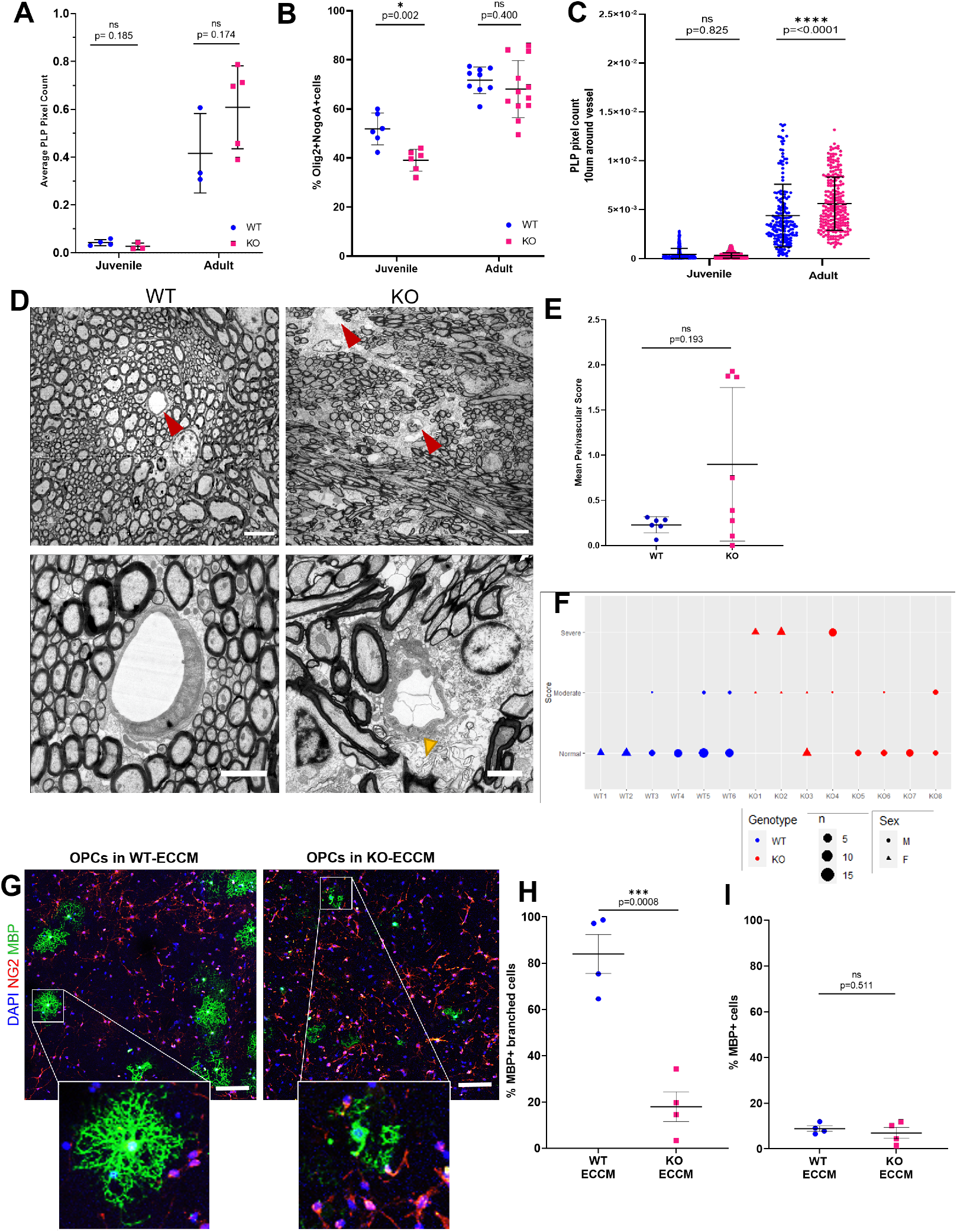
*Atp11b*KO white matter is abnormal. **(A)** There are no gross myelin defects as assessed by PLP+ pixel count in deep white matter of KO rats, as juveniles (t-test, p=0.185, t=1.536, df=5) or adults (t-test, p=0.174, t=1.540, df=6) (each point represents the average pixel count from six images of the deep white matter of one animal). **(B)** Significantly fewer NogoA+ oligodendroglia are present in the juvenile KO compared to WT animals (t-test, p=0.002, t=3.982, df=10), but not at the adult age (t-test, p= 0.400, t=0.8605, df=19) **(C)** In adult KO animals more PLP+ labelling is present in a 10µm wide ring adjacent to small vessels, compared to WT rats (One-way ANOVA, Tukey’s multiple comparison test, Juvenile: adjusted p=0.825, q=1.217, df=1314, Adult: adjusted p= <0.0001, q=10.28, df=1314) (each point represents the pixel count for a single blood vessel, accounting for area and adjusted across experiments for the overall PLP pixel count for each image.) **(D)** EM examples of perivascular changes in KO animals compared to WT (red arrowheads indicate blood vessels) at lower magnification (scale bar= 5µm) and at higher magnification (scale bar= 2µm) with yellow arrowhead indicating myelin debris **(E)** Quantified mean perivascular score per animal, (Mann-Whitney U =13.50, p=0.193). **(F)** Heterogeneity is seen between perivascular areas within animals; plotted as modal rater scores for each perivascular region analysed. **(H)** Oligodendrocyte precursor cells (OPCs) grown in endothelial cell conditioned media (ECCM) from cultures of KO-ECs show less branched morphology of MBP+ cells, compared to OPCs grown in ECCM from cultures of WT-ECs, as shown by immunostaining for oligodendroglial markers NG2 (red) and MBP (green), (scale bar 20µm) and percentage of branched MBP+ cells (t-test, p=0.0008, t=6.252, df=6) **(I)** but no difference in the number of MBP+ cells **(J)**, (t-test, p=0.511, t=0.6987, df=6). All graphs show mean +/- SD.

To assess whether this spatial and temporal correlation of EC dysfunction and oligodendroglia changes could be linked, we isolated these cell types from neonatal WT and KO animals for in vitro experiments. In the SHRSP, we previously showed that SHRSP ECs secrete more HSP90α than controls, leading to arrest of oligodendrocyte precursor cell (OPC) differentiation, reducing the number of mature myelin basic protein (MBP)-positive cells(22). Here, in a similar experiment, using cultured *Atp11b*KO ECs to condition media added to WT OPCs, there was no difference in the number of mature MBP+ oligodendroglia, but instead these *Atp11b*KO EC CM-treated MBP+ oligodendroglia showed a marked change in cell morphology with few and short processes, compared to the complexity of branching in controls, indicative of a maturation block at a later stage (**Fig. 3G-I**). WT oligodendroglia express *Atp11b* at the transcript(33) and at the protein level (**Suppl. Fig. 3A**), but isolated OPCs from neonatal *Atp11b*KO rats showed no difference in their survival, proliferation (Ki67+) or differentiation (MBP+) compared to WT OPCs in vitro (**Suppl. Fig. 3B-E**), indicating that the maturation block and morphological changes are an EC effect.

Thus, we have evidence of white matter damage selectively around smaller vessels that emerges as the animal ages, related to endothelial dysfunction in the *Atp11b*KO rat despite lack of hypertension. To relate this further to human SVD, we next assessed the rats with MRI, as diagnosis of SVD is in this way.

### The *Atp11b*KO rat has MR changes similar to human SVD

A greater extent of white matter changes as seen on MR correlates with a worse cognitive state(34) and allows SVD diagnosis. We performed longitudinal MR scanning using T1-weighted, T2-weighted, fluid attenuated inversion recovery (FLAIR) and T2*-weighted sequences on a cohort of rats both at 3-4 months and 9-10 months and found clear differences to the WT controls, with all KO rats having ventriculomegaly at the later age, suggestive of neurodegeneration and consistent with brain atrophy and ventricular enlargement in human SVD(35)(**Fig. 4A,B**). Spontaneous ventriculomegaly with age has previously been noted in adult Sprague-Dawley rats(36, 37) but here it is accelerated in *Atp11b*KO rats. Furthermore, there was also MR evidence of microbleeds in the KOs at both ages (**Fig. 4C,D**), again consistent with human SVD(38). The majority of apparent microbleeds in the older KOs were new, with only one lesion persisting from the younger age. However, there was no detectable change in white matter structural size measured at three locations in the corpus callosum, consistent with the focal white matter changes seen on pathology (**Suppl. Fig. 4**). We next analysed how these MR and pathological changes affected rat behaviour.

**Fig. 4.**
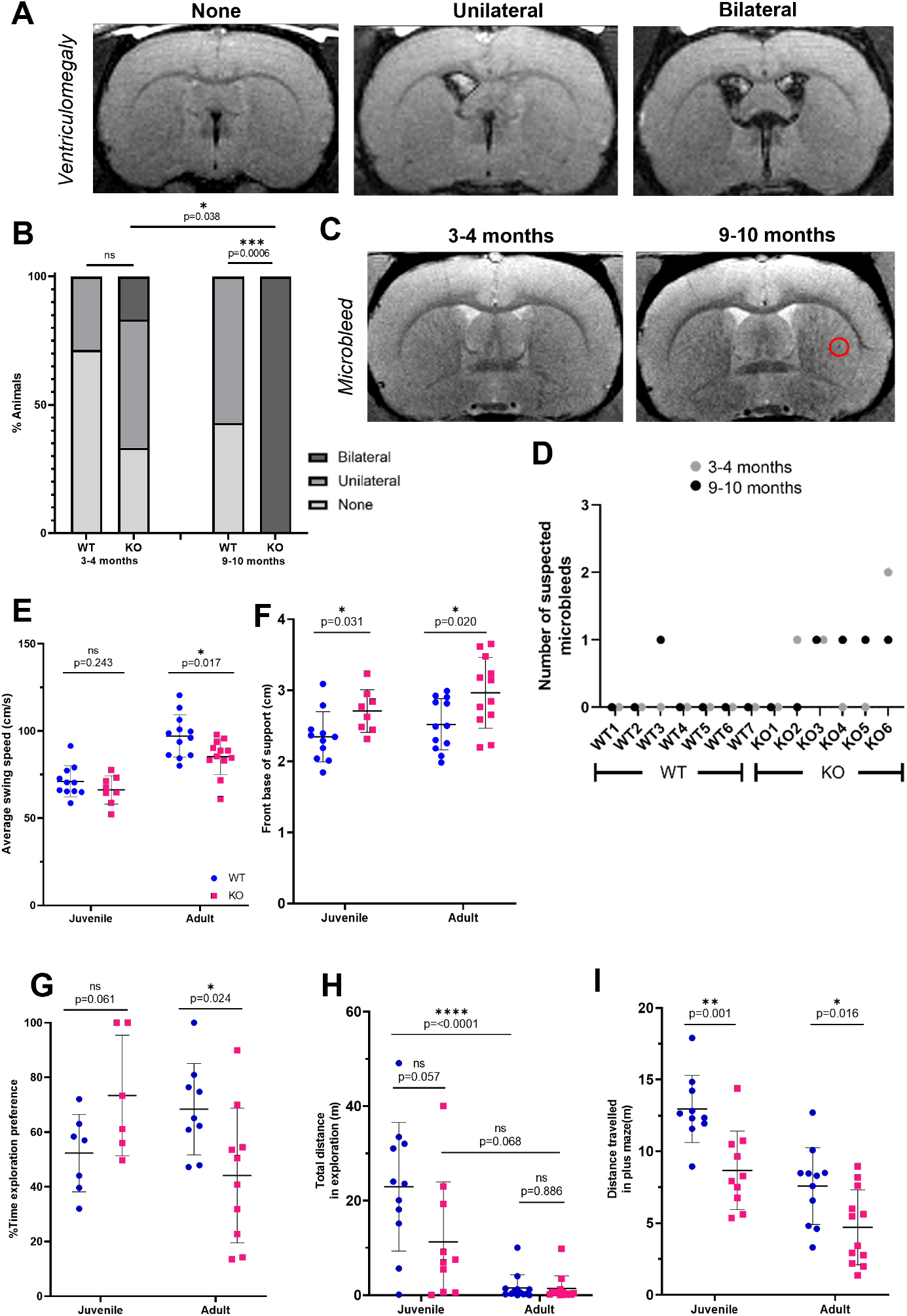
The *Atp11b*KO rat has MR and behavioural changes similar to SVD. **(A)** Illustrative images of the lateral and third ventricles on fluid-attenuated inversion recovery (FLAIR) sequences showing (from left to right) normal ventricles, unilateral ventricular enlargement and bilateral ventricular enlargement. **(B)** Significant difference in ventriculomegaly at 9 to 10 months of age between WT and KO (Fisher’s exact test, p= 0.0006). Significant difference in the relative proportions of normal, unilateral and bilateral scores between 3-4 month and 9-10 month old KO rat (Wilcoxon signed-rank test, p =0.038) but not in the WT (Wilcoxon signed-rank test, p =0.317). **(C)** Illustrative images of same rat brain at 3-4 months which then develops a suspected microbleed at 9-10 months (red circle).**(D)** KO animals have more suspected microbleeds at both ages compared to WT. **(E,F)** KO animals have altered gait shown through analysis using Catwalk, with **(E)** reduced average swing speed of all paws at the adult age (t-test, p=0.017, t=2.572, df=22) but not at the juvenile age (t-test, p=0.243, t=1.210, df=17) and **(F)** increased front base of support at both juvenile (t-test, p=0.031, t=2.348, df=17) and adult (t-test, p=0.020, t=2.499, df=22) ages. **(G,H)** Memory deficits emerge in KO animals - novel object recognition **(G)**, where KO animals spend less time investigating the novel object compared to WT at the adult age (t-test, p=0.024, t=2.482, df=17) suggesting memory failure, despite no difference in distance travelled compared to WT animals, at both ages **(H)** (t-tests, Juvenile: p=0.057, t=2.024, df=19; Adult: p=0.886, t=0.145, df=24). In addition, WT animals travel significantly less as they age (ANOVA, adjusted p=<0.0001, q=7.421, df=40), while KO animals show no significant difference, (ANOVA, adjusted p=0.068, q=3.600, df=40)(h) suggesting early deficits. **(I)** KO animals travel less distance in the plus maze compared to WT at both juvenile and adult stage, potentially indicating apathy. (t-tests, juvenile: p=0.001, t=3.762, df=18; adult: p=0.016, t=2.607, df=21).

### The *Atp11b*KO rat has behavioural changes similar to SVD

SVD patients can present with changes in altered gait, cognitive function and more subtle neuropsychiatric symptoms such as increased apathy(39), so we chose behavioural tests to probe these parameters. Similarly to humans with SVD, KO rats have problems with mobility, measured using the Catwalk equipment. *Atp11b*KO animals are slower (reduced swing speed (**Fig. 4E**)) and more unsteady (increased base of support (**Fig. 4F**)). They also have cognitive problems, with less interaction with novel objects in the Novel Object Recognition (NOR) test compared to WT age-matched controls in the adult age group, suggesting poorer memory that develops with age (**Fig. 4G**), despite normal movement in terms of distance travelled (**Fig. 4H**). Furthermore, in the elevated plus maze, KO rats at both ages travelled significantly less distance overall (**Fig. 4I**). This may reflect mobility issues, although KO rats move without problem for food and breeding, suggesting an additional degree of apathy, which was also identified in the human study(39).

## Discussion

Here, we show that the *Atp11b*KO rat has EC dysfunction, white matter pathological and MR-identified changes, and a behavioural phenotype similar to human SVD in the absence of hypertension and supporting the finding of genetic vulnerability to human sporadic SVD. This overturns the classical view that hypertension is required for the changes in the SHRSP and in human sporadic SVD, and refocuses our attention on intrinsic endothelial dysfunction as the key mechanism underpinning this disease, occurring early and inducing vulnerability of the white matter. This may explain why some SVD-affected individuals might be particularly sensitive to additional later exposures to extrinsic vascular risk factors such as hypertension and why it has proved difficult to prevent WMH progression even with intensive blood pressure reduction(3). We show that the loss of *Atp11b* is sufficient to cause EC dysfunction, molecularly and structurally, which alters with age and disease progression, and leads to disordered communication with surrounding oligodendroglia, laying the ground to identify therapeutic targets to reverse these changes. ATP11B is a phospholipid flippase, thought to move the phospholipids phosphatidylserine (PS) and phosphatidylethanolamine (PE) from the outer surface of a membrane to the inner, either at the plasma membrane or at vesicle membranes involved in vesicular transport(40). *Atp11b* deletion is therefore likely to alter membrane dynamics, with possible effects on intracellular transport (e.g. shuttling of endothelial NO to the EC surface), and on membrane receptors and ion channels, hence potentially altering signalling. Furthermore, exposed plasmalemmal PS and PE are known ‘eat me’ signals for myeloid cells(41). Here, we have a global rat KO of *Atp11b*, and so even though the EC-effect seems marked, other cells may also be affected, though there is potential compensation by other flippases (e.g. ATP11A or C, or others). There is impairment of hippocampal synaptic plasticity in a global-*Atp11b*KO mouse, but other phenotypes were not described(42), and so conditional mutants will be needed to unpick the relative contributions of each cell type to each pathology. Our previous identification of a SNP in *ATP11B* associated with human SVD MR changes indicates that the suggested effect on membranes has human relevance, and worth further investigation. We already know that the rare and familial SVD cathepsin A-related arteriopathy with strokes and leukoencephalopathy (CARASAL) also has a mechanism that links EC dysfunction to OPC maturation arrest and myelin abnormalities(43), suggesting that there may be other molecular deficits affecting this crosstalk that are hitherto unrecognised causes of sporadic SVD. We believe that focussing on these mechanisms, rather than hypertension, will be fruitful, especially as it is increasingly recognised that vascular changes contribute to several dementia types, including FTD and AD(44, 45). New European SVD guidelines lament the lack of effective treatments for SVD(12) and clearly state that although hypertension can contribute to the disease and should be treated, other therapies are needed. Here, we show that there is now a real opportunity to target EC dysfunction and its crosstalk with oligodendroglia for treatment of dementia.

## Materials and methods

### Study design paragraph

Our research objective is to understand small vessel disease by generating and characterising a normotensive rat transgenic for a relevant mutation. Sample sizes were calculated to give 80% power to detect a 30% outcome difference. We followed ARRIVE2 guidelines to minimize bias, with mixed-sex groups, and age-matching for WT and KO. Animals were only excluded from samples when culls were essential – such as injury from fighting.

Researchers were blinded to genotype during experiments for all blood pressure, behavioural and MRI data collection, and for image analysis, an ImageJ macro was used to anonymise and randomly order image titles for blind manual counting. Unless otherwise stated, for all graphs for immunofluorescence staining quantification each point represents a different animal/cell culture (biological replicate), calculated from a minimum of 5 fluorescence images (technical replicates) to generate one average value. For all graphs for western blot densitometry analysis, each point represents a different cell culture lysate (biological replicate), an average value calculated from three western blot experiments (technical replicates). For behavioural experiments, each point represents a different animal (biological replicate). Extreme outliers were defined as being 3xIQR outside the median value and were excluded from analysis.

### Generation of *Atp11b*KO rat

The rat *Atp11b* gene is located on rat chromosome 2 (GenBank accession number: XM_017591376.1; Ensembl: ENSRNOG00000052116) with the ATG start codon in exon 1 and TAG stop codon in exon 30. KO rat generation was designed by and purchased from Cyagen Biosciences Inc. The rat strain used was the standard Sprague-Dawley background rather than Wistar-Kyoto as in the SHRSP for ease of molecular genetics. Exons 5 to 8 were selected as target sites and guide RNAs chosen: gRNA1 (matches reverse strand of gene): ACAAAATGCTCACGCATCGAAGG gRNA2 (matches forward strand of gene): AGTGTAAAACTGT-CATCGGCTGG. Cas9 and gRNAs were injected into fertilised eggs for KO production and screened for deletion of this segment using PCR primers: Rat *Atp11b*-pair1-F: GGGTGCTCTAACTGCTCCAAGGTT Rat *Atp11b*-pair1-R: TAGATGCTGAAACAGACAAAAGCTACCAC PCR product size Rat *Atp11b*-pair1, giving a wildtype allele fragment of 7050 bp and a mutant allele fragment of 700 bp, thus deleting 6350 bp. Two F0 founders were generated, RatID#18 – male - missing 6352 bases on 1 allele and 6342 on other allele and Rat-ID#28 - female - missing 6349 bases on one allele only. These were bred with WT rats, leading to 6 F1 rats, four missing 6352 bases on 1 allele (from Rat-ID#18) and 2 missing 6349 on1 allele (from Rat-ID#28). These deletions lead to predicted truncated amino acid sequences which are identical MWRWVRQQLG FDPPHQSDTR TIYIANRFPQ NGLYT-PQKFI DNRIISSKYT VWNFVPKNLF EQFRRVANFY FLIIFLVQLM IDTPTSPITS GLPLFFVITV TAIKQTSGTR ESLASWSQIK KYKRNFWCGC IHWDGNKDGI ELQE-QVTEKI CCREINEHIF NNLSNNPYF CCREINEHIF NNLSNNPYF. Therefore, we considered these the same, and formed the KO rat by interbreeding the F1 generation. Off target analysis identified three potential off-target sites for gRNA1 and two for gRNA2, so Cyagen designed primers around these sites, and generated and sequenced PCR amplicons but found no mutations (data available on request).

### Housing and husbandry

All work was carried out under UK Home Office project licence PADF15B79. *Atp11b*KO animals were bred as heterozygous breeding pairs to generate WT and KO littermates for in vivo work. For neonatal cell isolation, WT/WT and KO/KO breeding pairs were used to ensure correct genotype at birth. Heterozygote animals were used in outbreeding but not used as part of study.

### Termination

For OPC isolation, pups at postnatal day 0-2 were terminated by an overdose of Euthatal (150mg/kg; Merial) administered intraperitoneally. For brain microvascular endothelial cells, (BMECs) pups at postnatal day 5-7 were terminated by a rising concentration of carbon dioxide (C_O_2). For extraction of protein lysates where fresh frozen tissue was required, animals were terminated by an overdose of Euthatal (150mg/kg) and brains extracted and stored in PBS temporarily until processed. For immunohistochemistry (IHC), animals were first anaesthetized with Isoflurane by inhalation, then by intraperitoneal injection with Ketamine (Vetalar®) and Medetomidine (Domitor®). Intracardiac perfusion fixation was performed, first with phosphate buffered saline (PBS) then with 4% paraformaldehyde (PFA; Sigma-Aldrich; diluted in PBS (Gibco)).

### Blood pressure measurement

Blood pressure was measured weekly using the CODA™ Non-Invasive Blood Pressure System, and body weight was also recorded. Animals were habituated to the tube in advance to avoid stress. For each weekly reading, five test cycles were carried out before 20 recorded cycles for each animal. Readings were excluded by the system if they did not fit the expected curve. A weekly blood pressure average for each animal was obtained from a minimum of 8 recorded cycles. Experiments were carried out by a researcher blinded to genotype. Raw data was extracted from the operating systems by a different researcher and processed.

### Open Field and Elevated maze

Animals were handled prior to testing to minimize stress. The protocol was carried out by an independent researcher, who was blind to genotype to prevent experimenter bias. Open field or elevated maze was set up directly above the camera and AnyMaze software used to record when animals entered the centre of the field and the time spent in the edges. The animal was initially placed in the corner facing the wall for open field and placed in middle of the elevated maze. The open field test was run for 20 minutes, and the elevated maze test for 5 minutes, with the researcher hidden.

### Catwalk

Experiments were based on published methods with minor modifications(46, 47). The automated gait analysis apparatus (Noldus, Wageningen, Netherland) consists of a glass plate walkway equipped with light-emitting diodes (LEDs). Light from the LEDs is emitted inside the glass plate, internally reflected, and refracted to the glass plate except in the area where the animal’s paw makes contact with the plate. A high-speed colour camera positioned underneath the glass plate captures images of the illuminated area and sends the imaging information to a computer that runs the analysis system (CatWalk XT version 10.6 software; Noldus). Animals walked on the glass plate walkway 3 times on the day before evaluation to habituate them. A satisfactory measured walk was defined as an uninterrupted walk across the walkway that left >3 footprints for each hind limb. Recordings were repeated until three satisfactory walks were obtained.

### Novel Object Recognition

Animals were handled prior to testing to minimize stress. Protocol was carried out by an independent researcher, who was blind to genotype to prevent experimenter bias. Experiments were based on published methods with minor modifications(48). The test area consisted of an open field set up directly below the camera and AnyMaze software used to record when animals entered areas surrounding objects. After habituation to both the test area and familiarisation to two identical objects (two lego towers), a new object (lightbulb) replaced one of the original objects and the animal was initially placed in the corner facing the wall of the open field when recording was started. The percentage time exploration preference was calculated as a ratio: time spent exploring the novel object/time spent exploring the familiar object.

### Magnetic Resonance Imaging Acquisition and Processing

*In vivo* MRI data was acquired in a longitudinal way on age-matched *Atp11b*KO and Sprague-Dawley rats at both 3-4 months and at 9-10 months of age. Anaesthesia was induced in an adapted chamber with 4% isoflurane and a 30:70 O_2_/N_2_O ratio. Animals were then transferred to the MRI instrument cradle. A hot air blower was used to regulate physiological temperature (37±1°C). The head was secured laterally by conical ear rods and longitudinally by the nose cone used for anaesthetic gas delivery. The animals breathed spontaneously through a facemask, with isoflurane delivered at a constant flow mixed with a 40:60 ratio of O_2_/N_2_O (1 L min-1). Isoflurane concentration varied (1.5-3%) to maintain stable respiration rates (40-100 bpm). Respiration was monitored using a pressure sensor connected to an air-filled balloon placed under the animal abdomen. On completion of imaging, isoflurane delivery was stopped and animals observed until full recovery.

Images were acquired in a 400-mT/m gradient set 7T Bruker scanner (Bruker Medical GmbH, Germany) with an 86-mm ID quadrature radiofrequency coil for transmission and a 4-channel phased array coil for reception. The acquisition parameters are listed in the table below. A localizer scan was performed to ensure correct positioning before T_1_-weighted (T1W), T_2_-weighted (T2W), T_2_^*^-weighted (T2^*^W) and T2W with fluid attenuated inversion recovery (FLAIR) images were acquired from the caudal to cranial cerebral cortex. Images were uploaded to Carestream Vue PACS (Carestream Health, Onex, Canada) for quantitative and qualitative image assessment.

Quantitative assessment was performed on T2W images using the graphics line measurement tool. Corpus callosum was measured in the coronal plane at three locations of one slice rostral to the merging of the anterior parts of the anterior commissure. Qualitative assessments for ventriculomegaly and potential microbleeds were performed using a combination of the four sequences acquired with particular emphasis on T2*W for microbleeds and FLAIR for ventriculomegaly. These methods were adapted from our human-based methods(49).

| Protocol | Parameters | Resolution |
| --- | --- | --- |
| T1W RARE | FOV 30x30mm, Matrix 256x256x30, SLT 0.5mm, TE 11.3ms, TR 1200ms | 0.12x0.12x0.50mm |
| T2W RARE | FOV 30x30mm, Matrix 256x256x30, SLT 0.5mm, TE 45ms, TR 2500ms | 0.12x0.12x0.50mm |
| T2*W FLASH | FOV 30x30mm, Matrix 256x256x30, SLT 0.5mm, TE 10ms, TR 700ms | 0.12x0.12x0.50mm |
| T2W FLAIR | FOV 30x30mm, Matrix 192x192x15, SLT 1.0mm, TE 45ms, TR 8000ms, TI 1860ms | 0.16x0.16x1.0mm |

FOV: field-of-view; SLT: slice thickness; TE: echo time; TR: repetition time; RARE: rapid acquisition with relaxation enhancement; FLASH: fast low angle shot; TI: inversion time

### Preparation of brain endothelial cells

As adapted from a previously published method(50). Brains were extracted from rats aged postnatal day 5-7 and placed in working buffer (Hank’s balanced salt solution (HBSS) (Life Technologies) with 0.5% chromatographically purified bovine serum albumin (BSA) (First Link UK), 0.5% HEPES (Gibco), 0.5% pen/strep). Meninges and cerebellum were removed and brains were homogenised in 5ml working buffer with a Dounce tissue grinder. Homogenate was isolated from the buffer by centrifugation for 5 minutes at 1800 revolutions per minute (RPM) at 4^°^C. Blood vessel fragments were pelleted from this homogenate by centrifugation in 22% BSA for 15 minutes at 3000 RPM at 4^°^C. The pellet of blood vessel fragments was kept on ice in working buffer while the remainder was centrifuged again, for a total of 5 spins. Blood vessel fragments were pipetted onto an upside down 70µm cell strainer to remove single cells then washed into a pooled mixture with working buffer. This was spun down (5 minutes at 1800 RMP, 4^°^C) and the pellet then digested in 10ml collagenase/dispase (1mg/ml; Roche) and DNase I type IV (40µg/ml; Sigma-Aldrich) for 45 minutes at 37^°^C with regular agitation.

The resultant dissociated endothelial cells were washed in working buffer and plated at 500,000 cells/well on collagen IV and fibronectin (100µg/ml and 50µg/ml respectively; Sigma-Aldrich; overnight, 4^°^C) coated coverslips in a 24-well plate in 1ml Endothelial Growth Media-2 Bullet Kit (EGM-2; Lonza; Endothelial basal media-2 with 2% Fetal Bovine Serum (FBS), 0.4% hFGF-beta, 0.1% hEGF, 0.1% VEGF, 0.1% R3-IGF-1, 0.1% Ascorbic acid, 0.1% Gentamycin/Amphotericin-B, 0.1% puromycin (Sigma-Aldrich; 5mg/ml in Dulbecco’s phosphate buffered saline (DPBS; Sigma-Aldrich)) and 0.04% hydrocortisone). Cells were grown in 7.5% carbon dioxide (CO_2_) and medium was changed every 2-3 days. Cultures were used in experiments once they had regions of confluence (before 10 days).

### Preparation of OPCs

As adapted from a previously published method(51). Brains were extracted from rats aged postnatal day 0-2 and cortices isolated in MEM. Meninges were removed then cortices were minced with fine scissors and digested in MEM with papain (1.2U/ml; Worthington), L-cysteine (0.24mg/ml; Sigma-Aldrich) and DNase I type IV (40µg/ml) for 1 hour at 37^°^C. Cells were diluted in culture medium (Dulbecco’s modified Eagle medium (DMEM; Gibco) with 10% FBS (Gibco), 1% pen/strep), spun down (5 minutes at 1000 RPM), then resuspended with a 1ml pipette. Cells were resuspended in culture medium (1.5 brains in 10ml per flask) and grown in vented T75 flasks coated with poly-D-lysine (PDL; Sigma-Aldrich; 5µg/ml in water; minimum 1 hour coating, 37^°^C) in 7.5% CO_2_.

10-14 days after dissection, the vented cap of the flasks was sealed and the flasks were placed on an orbital shaker at 240 RPM at 37^°^C. After 1 hour the medium was removed (containing loosely attached microglia), fresh medium added, and the flask returned to the shaker for 16-18 hours. This medium was removed and plated on 10cm plastic petri dishes for 20-25 minutes (to remove remaining microglia) after which the cells were spun down (5 minutes at 1000 RPM). Cells were resuspended in 5ml culture medium, gently triturated through a 21 gauge needle, counted, and plated at a density of 50,000 cells in a single droplet of SATO medium (DMEM with 1% pen/strep. 1% ITS supplement (Sigma-Aldrich), 16µg/ml putrescine (Sigma-Aldrich), 400ng/ml L-thyroxine (Sigma-Aldrich), 400ng/ml Tri-iodothyroxine (Sigma-Aldrich), 60ng/ml progesterone (Sigma-Aldrich), 100µg/ml BSA (fraction V) (Sigma-Aldrich)) per PDL coated coverslip in a 24 well plate, topped up to 0.5ml SATO medium per well after 20 minutes at 37^°^C for cells to stick down. OPCs were grown for 2 days and the medium was not changed during this time.

### Conditioned media experiments

Endothelial cells (HUVECs and BMECs) were grown in their normal culture medium until reaching confluence and forming tight junctions (∼10 days *in vitro* (DIV) for BMECs, 5 DIV for HUVECs). At this point the media was changed and this was conditioned for 2-3 days. Media was then removed and stored at -20^°^C until required. When defrosted, conditioned media was mixed at a 1:1 ratio with SATO media. Cells grown in conditioned media were prepared as normal. Conditioned media was added to OPCs after the step at which they were left for 20 minutes to stick down. Cells were left for 3 days in conditioned media before being fixed and analysed.

### Immunofluorescence

Brains were extracted from animals after perfusion fixation and incubated for no more than 4 hours in 4% PFA at 4^°^C. PFA-fixed brains were then incubated first in 15% sucrose (overnight, 4^°^C; Sigma-Aldrich) and then in 30% sucrose (overnight, 4^°^C). Sections were cut at 10µm thickness using a cryostat and collected in series so that each slide for immunostaining had a spread of coronal sections of the deep white matter.

Eyes were extracted following perfusion and fixation and washed with PBS before storage in 30% w/v sucrose. Eyes were dissected using an adapted version of a previously published method(28). Immunofluorescence staining was performed with the retina still attached to the sclera of the back of the eye due to tissue fragility. Following staining, the retina was quartered and the sclera discarded before flat-mounting for imaging. These were imaged with a Zeiss wide-field microscope.

### For brain and cell immunofluorescence

Where necessary for antibodies, antigen retrieval was carried out by boiling sections for 10 minutes in a microwave in citrate buffer (Vector). Sections were then cooled for 20 minutes in running tap water. Sections were blocked for at least 1.5 hours at room temperature in blocking solution (10% heat inactivated horse serum (HIHS), 0.5% triton (Fisher) in PBS). Primary antibodies, diluted in blocking solution, were added and the sections were incubated at 4^°^C overnight in a humid chamber. The sections were washed in PBS then incubated with secondary antibodies, diluted in blocking solution, for 1.5 hours at room temperature. Sections were washed in PBS before adding DAPI (VWR) for <1 minute. Sections were mounted with Fluoromount (Southern Biotech). Fluorescent images were captured using Leica SP8 confocal microscope system.

### For retinal immunofluorescence

Dissected eyes were first incubated with a blocking solution with 10% Serum and 0.05% Triton-X in PBS before overnight incubation with biotinylated *Griffonia simplicifolia* isolectin B4 mixed at a 1:500 dilution in blocking solution. Samples were washed three times for 5 minutes with PBS before incubation with streptavidin with fluorescent tag (Alexa Fluor 647 – Thermofisher) in blocking solution at 1:1000 dilution for 1.5hrs. Samples were washed again before incubation with Hoechst for 1 minute. Samples were rinsed with PBS before retinal removal and mounting. Retinal flat-mounts were imaged with a Zeiss wide-field microscope, generating a tiled scan of the retina.

### Electron microscopy

Procedure carried out by an independent researcher who was blind to genotypes. Rats were transcardially perfused; first with PBS (< 2 min), then with fixative (4% paraformaldehyde w/v, Sigma; 0.1-2 glutaraldehyde, v/v, Electron Microscopy Sciences (EMS) or TAAB Laboratories Equipment Ltd.; in 0.1 M phosphate buffer, Sigma; 10 min). The brains were extracted, post-fixed in the same fixative (24h at 4^°^C), and vibratome-sectioned (50 µm thick). Sections were post-fixed with 1% osmium tetroxide (EMS; in 0.1 M phosphate buffer; 30 min) then dehydrated in an ascending series of ethanol dilutions [Sigma; 50%, 70% (with 1% uranyl acetate; w/v; EMS), 95% and 100%] and acetone (EMS). Sections were then lifted into resin (Durcupan ACM; EMS), left overnight at room temperature, then placed on microscope slides, covered with coverslips, and resin cured at 65^°^C for 3 days.

Subcortical white matter (internal capsule) was imaged with brightfield light microscopy then cut from the slides, mounted on plastic blocks (Ted Pella) and ultrathin sectioned (70 nm thick) for electron microscopy. Serial sections were collected on to single-slot, formvar-coated copper grids (EMS), contrasted with lead citrate (EMS) and Uranyless (EMS) and imaged with electron microscope (JOEL TEM-1400 plus). Magnifications were chosen to visualise each vessel with surrounding perivascular region for assessment.

### Image processing and quantification

#### For brain images

Cell counting was carried out manually using max projections of z-stacks of 10µm tissue at 20x generated with ImageJ and the Cell Counter plugin. For cultured endothelial cell CLDN5 quantification, cells were classified as CLDN5+ if the border with all neighbouring cells (delineated with ZO-1 staining) had CLDN5 present.

#### For retinal images

In the plexus, regions of interest (ROIs) were selected in ImageJ using the diameter of the optic disc as a relative measurement. ROIs were taken from both proximal and distal selections in each of the four cut sections of the retina. ROIs then underwent segmentation in FIJI to generate a binary representation of the vasculature. ROIs were inverted, contrast enhanced and segmented using Phansalkar auto local thresholding to binarise the image(52), images were then filtered(53) and staining abnormalities accounted for(54). Binarised ROIs then underwent skeletonisation via Voronoi tessellation as previously described(55). Radius data for the skeleton was measured during skeletonisation by defining pixels on the boundary of the binary mask and measuring the distance erased to produce skeletons. Vessel tortuosity and branching were measured using a Python script which defines branching points as vertices in the skeleton with greater than 2 neighbouring vertices. This allows quantification of tortuosity by measuring actual versus Euclidean distances between these points. Branching index was calculated as the number of branching points per µm^2^. Major vessels were split into arterioles and venules for analysis and average vessel diameter, tortuosity and branching indices measured manually in ImageJ. Average vessel diameter was measured by taking manual measurements across the vessel at 200µm intervals from vessel origin to terminal bifurcation.

#### For PLP pixel count analysis

Vessels were manually delineated using IsoB4 staining from max projections of z-stacks of 10µm tissue at 20x generated with ImageJ, and a concentric ring (10µm width) automatically generated around each vessel, adjusted to ensure that no two areas are analysed twice; a part of an outermost ring was subtracted if it overlapped with another vessel’s innermost ring.

### Statistics

Described within figure legends for each graph, all of which show mean +/- SD. Graphs and statistical calculations were made using GraphPad Prism version 9.0.0 for Windows, GraphPad Software, San Diego, California USA, www.graphpad.com or RStudio Team (2021) (RStudio: Integrated Development Environment for R. RStudio, PBC, Boston, MA, www.rstudio.com). When comparing between two groups, (typically WT and KO), t-tests (parametric data) or Mann-Whitney U test (non-parametric) were used. When comparing interactions of age as well as genotype, ANOVA was used. For MRI analysis a Fisher’s exact test was performed to analyse differences in ventriculomegaly and a Mann Whitney-U test for differences in corpus callosum size. Statistical analysis of ventriculomegaly proportions within a genotype longitudinally was performed using a Wilcoxon signed rank test. Significance codes were generated by GraphPad to report three significant digits. p<0.05 was set as criteria for statistical significance for all tests. P values less than 0.001 are summarized with three asterisks, P values less than 0.0001 summarized with four asterisks.

## Acknowledgments

Thanks to Ross Lennen for MRI scanning, Tetiana Poliakova for preliminary MRI analysis, Stephen Mitchell for help with EM and James Ashmore for his input with the statistics.

## Funding

UK Dementia Research Institute as UK DRI which was funded by the MRC, Alzheimer’s Society and Alzheimer’s Research UK (JW)

British Heart Foundation (AW) Fondation Leducq (JW)

Centre for Cognitive Ageing and Cognitive Epidemiology pilot fund (AW, JW)

Wellcome Trust Tissue Repair PhD fellowship (SQ)

Wellcome Trust Edinburgh Clinical Academic Trainee – Veterinary Sciences (TP).

## Author contributions

Most experiments: SQ, TP, JM

MRI experiments: AV, MJ

MR analysis: TP, AV, MJ, JW, ZT

Retinal analysis: AL, YG, MB.

PLP analysis: MW

Oligodendrocyte analysis: SB

Behavioural tests: MM

Blood pressure measurements: AO

Rat perfusion and care: AO, WM

Conceptualization and direction: JW and AW

Writing – original draft: SQ, AW

Writing – review & editing: all authors

## Data and materials availability

All data are available in the main text or the supplementary materials. Code is available at https://github.com/mobernabeu/normotensiveSVDrat for retinal vessel analysis and PLP pixel count.

## Supplementary Figures

**Fig. S1.**
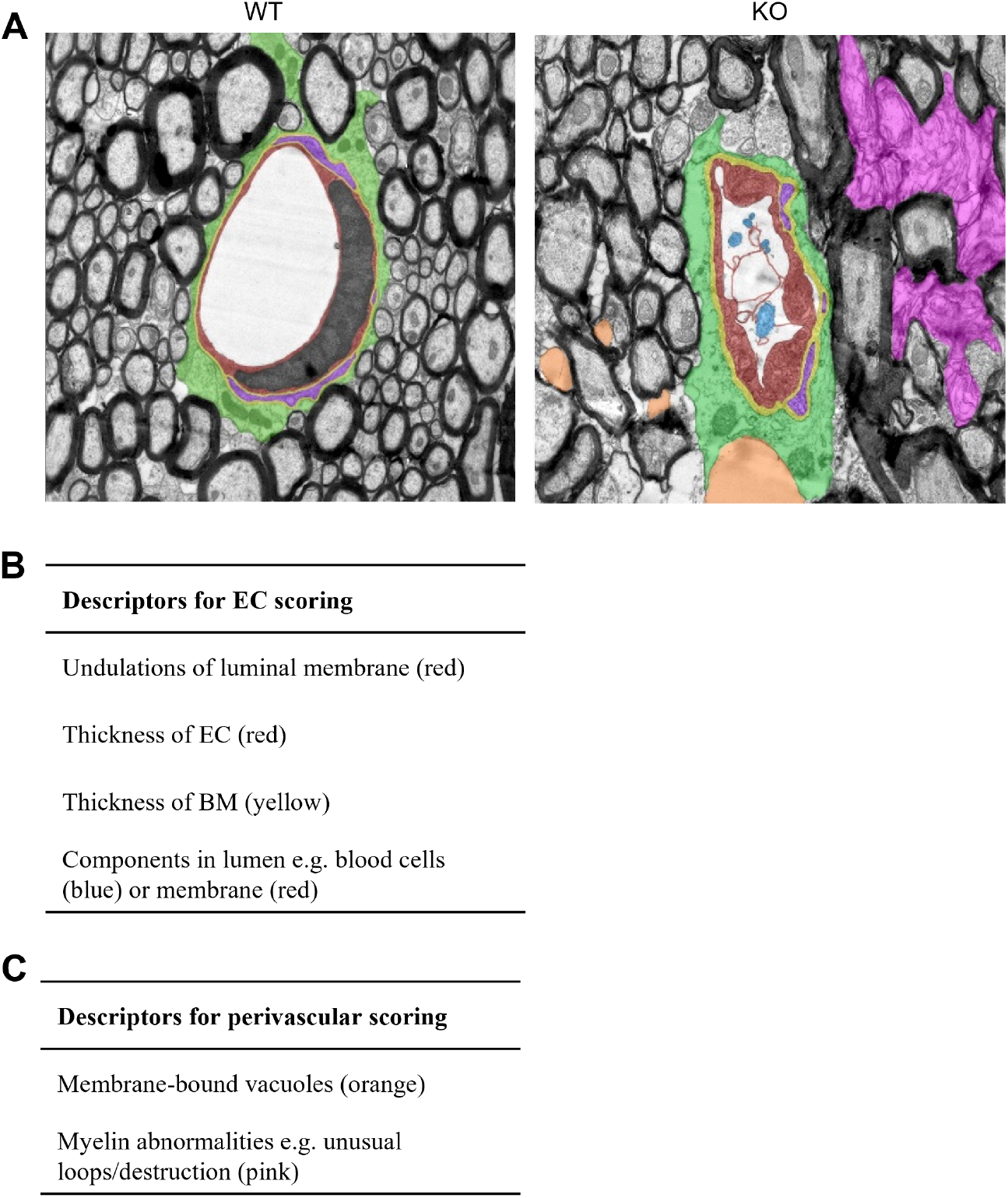
Electron microscopy pathological grading system. **(A)** Example of normal and abnormal images, with tables of descriptors **(B,C)**. Additional features, not scored: astrocytes, green; pericytes, purple; EC nuclei, grey.

**Fig. S2.**
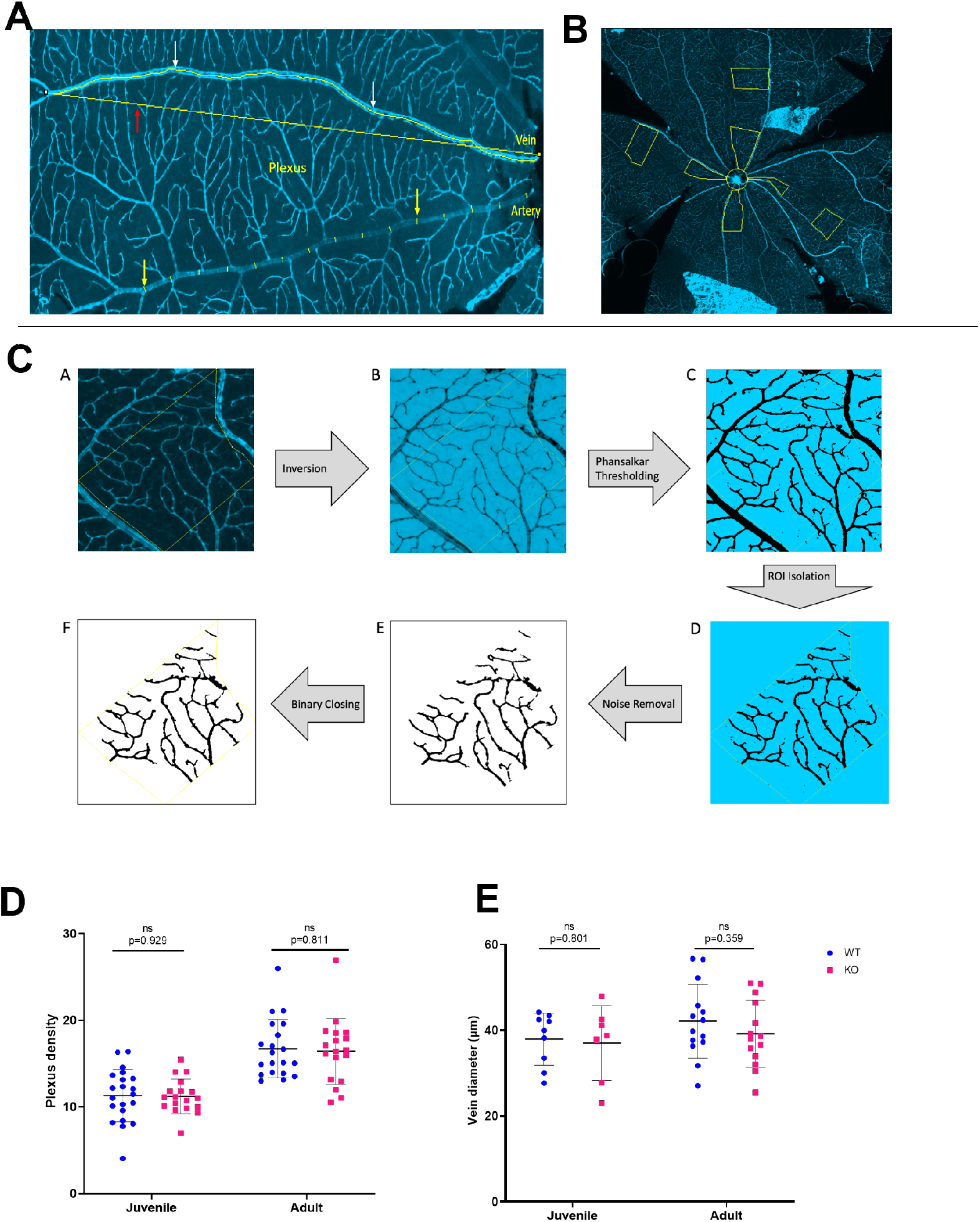
Retinal image ROI selection and additional data. **(A)** Representative image of tortuosity and diameter measurement in the major vessels. White arrows indicate measurement of total vessel length, red arrow indicates measurement of the Euclidean distance between vessel endpoints, yellow arrows indicate diameter measurements. **(B)** Representative image of ROI selection. Four proximal ROIs are located centrally, with three distal ROIs outside. A central ring is shown for measurement of the optic disc diameter. **(C)** Workflow with representative images of a distal ROI at various stages of processing to prepare for analysis. **(D)** Retinal plexus density is not significantly different between WT and KO animals at either juvenile age (t-test, p=0.929, t=0.089, df=37) or adult age (t-test, p=0.811, t=0.24, df=36). **(E)** Vein diameter is not significantly different between WT and KO animals at either juvenile age (t-test, p=0.801, t=0.357, df=14) or adult age (t-test, p=0.359, t=0.93, df=26).

**Fig. S3.**
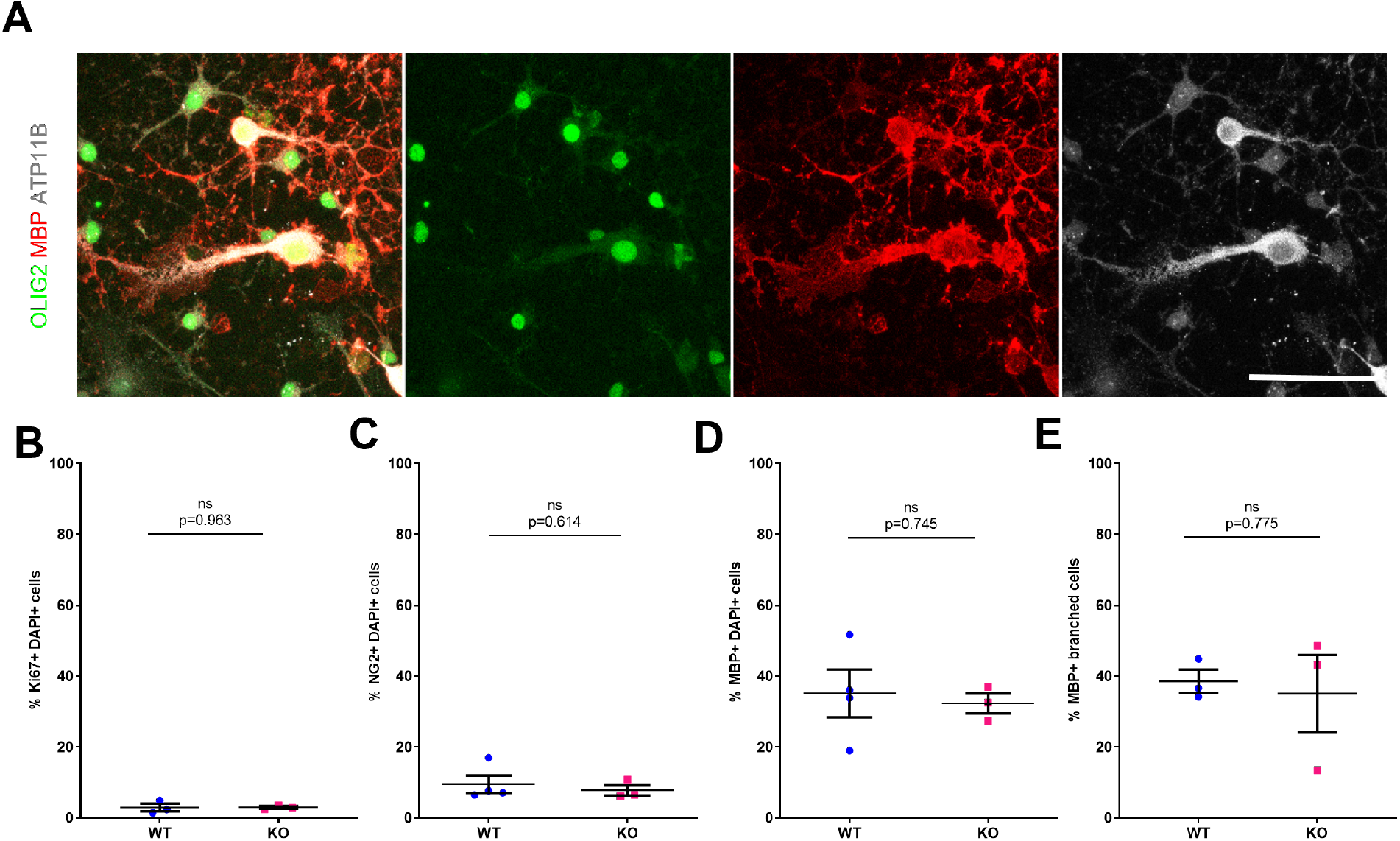
ATP11B expression in oligodendroglia *in vitro*. **(A)** ATP11B is expressed by WT OPCs in culture by immunofluorescence, scale bar 20µm, nuclear OLIG2 in green, MBP in red and ATP11B in grey. Loss of *Atp11b* does not have an effect on OPCs in vitro either in **(B)** number of Ki67+ proliferating cells (t-test, p=0.963, t=0.04888, df=4) **(C)** number of immature NG2+ cells (t-test, p=0.614 t=0.5380, df=5) **(D)** number of mature MBP+ cells (t-test, p=0.745, t=0.3442, df=5) or **(E)** number of branched MBP+ cells (t-test, p=0.775, t=0.3051, df=4).

**Fig. S4.**
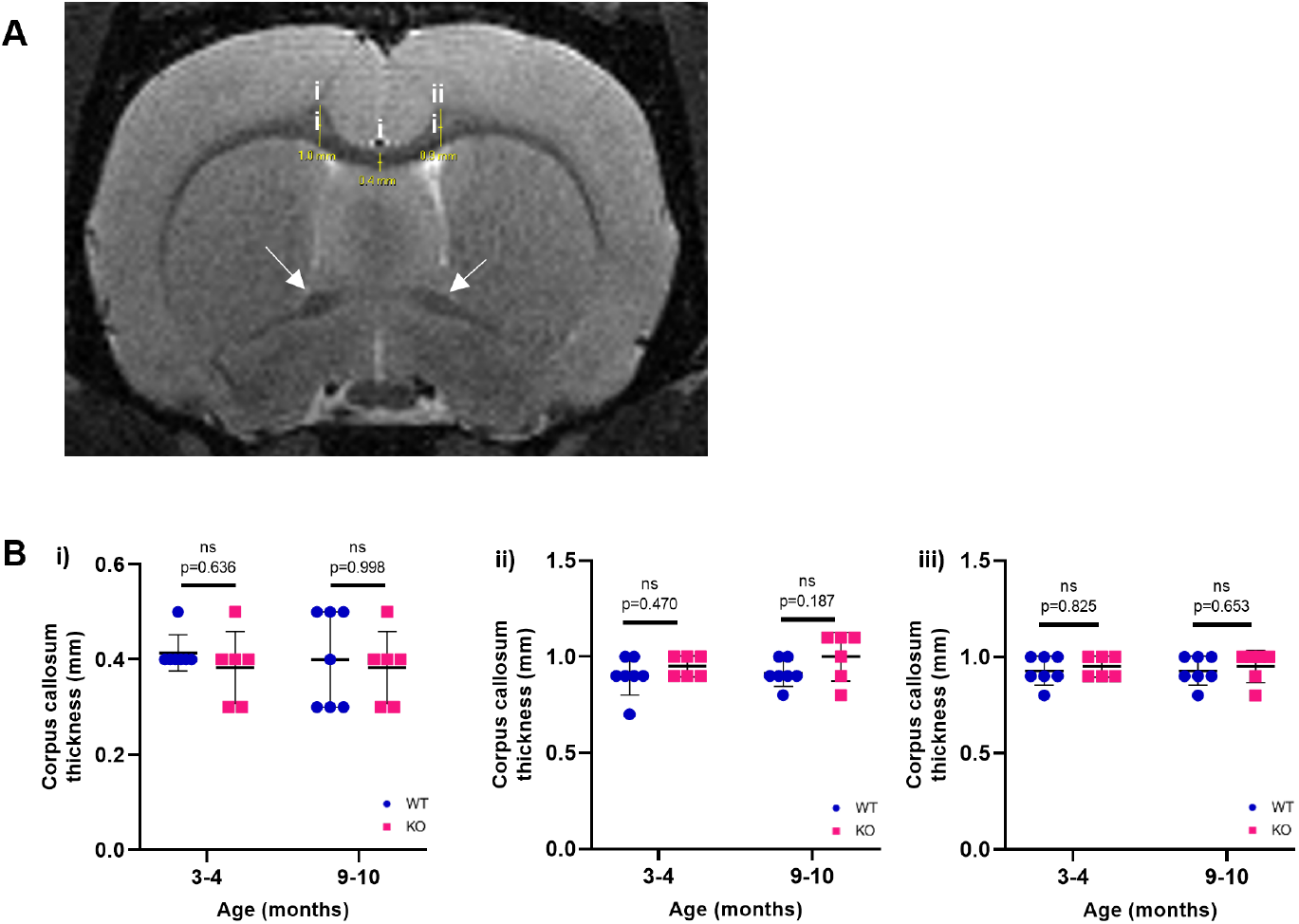
No change in *ATP11B*KO white matter size measurements by MRI. **(A)** Diagram demonstrating corpus callosum measurements on T2-weighted images taken in transverse section just prior to the anterior part of the anterior commissure merging (arrows). Three measurements were taken in positions i, ii and iii of the corpus callosum. **(B)** No significant difference in corpus callosum thickness between WT and KO at position i) (3-4 months: Mann-Whitney U =15.50, p=0.636 9-10 months: Mann-Whitney U =19, p=0.998; ii) (3-4 months: Mann-Whitney U =15, p=0.470; 9-10 months: Mann-Whitney U =11.50, p=0.187); or iii) (3-4 months: Mann-Whitney U =18, p=0.825; 9-10 months: Mann-Whitney U =17, p=0.653).

